# Rapid simultaneous acquisition of macromolecular tissue volume, susceptibility, and relaxometry maps

**DOI:** 10.1101/2021.06.08.447552

**Authors:** Fang Frank Yu, Susie Yi Huang, Thomas Witzel, Ashwin Kumar, Congyu Liao, Tanguy Duval, Julien Cohen-Adad, Berkin Bilgic

**Affiliations:** Radiology, University of Texas Southwestern Medical Center, Dallas, TX, United States; Department of Radiology, Harvard Medical School, Boston, MA, United States; Harvard-MIT Health Sciences and Technology, Massachusetts Institute of Technology, Cambridge, MA, United States; Athinoula A. Martinos Center for Biomedical Imaging, Charlestown, MA, United States; Q Bio, *Inc*., Redwood City, CA, United States; Vanderbilt University, Nashville, TN, United States; Radiological Sciences Laboratory, Stanford Medicine, Stanford, CA, United States; Institute of Biomedical Engineering, Ecole Polytechnique de Montreal, Montreal, QC, Canada

**Keywords:** Quantitative susceptibility mapping, myelin sensitive imaging, parallel imaging, multi-parametric imaging, MRI

## Abstract

**Purpose:** A major obstacle to the clinical implementation of quantitative MR is the lengthy acquisition time required to derive multi-contrast parametric maps. We sought to reduce the acquisition time for quantitative susceptibility mapping (QSM) and macromolecular tissue volume (MTV) by acquiring both contrasts simultaneously by leveraging their redundancies. The Joint Virtual Coil concept with generalized autocalibrating partially parallel acquisitions (JVC-GRAPPA) was applied to reduce acquisition time further.

**Methods:** Three adult volunteers were imaged on a 3T scanner using a multi-echo 3D GRE sequence acquired at three head orientations. MTV, QSM, R2*, T1, and proton density maps were reconstructed. The same sequence (GRAPPA R=4) was performed in subject #1 with a single head orientation for comparison. Fully sampled data was acquired in subject #2, from which retrospective undersampling was performed (R=6 GRAPPA and R=9 JVC-GRAPPA). Prospective undersampling was performed in subject #3 (R=6 GRAPPA and R=9 JVC-GRAPPA) using gradient blips to shift k-space sampling in later echoes.

**Results:** Subject #1’s multi-orientation and single-orientation MTV maps were not significantly different based on RMSE. For subject #2, the retrospectively undersampled JVC-GRAPPA and GRAPPA generated similar results as fully sampled data. This approach was validated with the prospectively undersampled images in subject #3. Using QSM, R2*, and MTV, the contributions of myelin and iron content to susceptibility was estimated.

**Conclusion:** We have developed a novel strategy to simultaneously acquire data for the reconstruction of five intrinsically co-registered 1-mm isotropic resolution multi-parametric maps, with a scan time of 6 minutes using JVC-GRAPPA.

## 1. Introduction

Quantitative susceptibility mapping (QSM) has emerged as a promising technique for measuring magnetic susceptibility, which has been shown to correlate with paramagnetic substrates in the brain, including iron which is critical for cellular function.^1,2^ Unfortunately, deriving susceptibility from phase data requires solving an ill-conditioned linear system, which leads to image artifacts. To address this shortcoming, it becomes necessary to impose spatial regularization^3–5^ or acquire additional volumes at different head orientations (COSMOS), the latter which has been shown to generate higher quality susceptibility calculations.^6^

Myelin, which is comprised of a lipid and protein bilayer, forms sheaths around axons and facilitates neural conduction. Myelin content can be probed using T_2_-relaxometry^7^, quantitative magnetization transfer (qMT)^8^, macromolecular tissue volume (MTV)^9^, and direct visualization of short transverse relaxation time component^10^. MTV estimates macromolecule content, and is highly correlated with qMT and T_2_-relaxometry.^9,11^ Importantly, MTV offers a straightforward acquisition with high spatial resolution and signal-to-noise ratio (SNR)^12,13^.

The information conferred through QSM and MTV could have important clinical applications in the management of neurological disorders that perturb myelin and iron content.^9,14–16^ Although both are acquired using GRE sequences, they are traditionally obtained separately. Unfortunately, this can lead to lengthy scan times and image mis-registration, complicating quantitative analyses. Herein, we exploit inherent redundancies between these two sequences by proposing a novel strategy to acquire whole-brain multi-orientation QSM and MTV simultaneously within the scan time of only one contrast. Additional quantitative parametric maps can be generated from the acquired data, including T_1_, R_2_*, and PD. Furthermore, we show that these multi-parametric maps can be leveraged to produce myelin and iron-sensitive susceptibility maps.

To enhance clinical feasibility, we propose the use of Joint Virtual Coil generalized autocalibrating partially parallel acquisitions (JVC-GRAPPA).^17^ Parallel imaging techniques such as GRAPPA allow for reduced image acquisition times.^18 19^ The weighting coefficients (GRAPPA kernels) are derived from a fully-sampled autocalibration signal (ACS) region. Unfortunately, acceleration factors higher than R=3 along one phase-encoding axis are generally avoided due to SNR degradation.^20^ To address this limitation, additional virtual coils (VC) can be generated using the complex conjugate symmetric k-space signals from the actual coils. The conjugate phase information provided by the VCs improves reconstruction and image quality.^19,21,22^

JVC-GRAPPA expands upon this by jointly reconstructing multiple echoes within the GRAPPA framework. In effect, a particular channel receives contributions from all image contrasts across all channels, while the phase information is converted into additional spatial encoding.^19,21^ We hypothesized that JVC-GRAPPA would yield image quality comparable to GRAPPA reconstructions while significantly reducing scan time.

The goals of our study were to 1) develop an acquisition scheme for collection of data suitable for reconstruction of QSM and MTV, reducing scan time to that of a single contrast, and 2) further reduce scan time using JVC-GRAPPA. If successful, our strategy could facilitate quantitative evaluation of neurological diseases.

## 2. Methods

MTV is acquired using a multi-flip angle (FA) 3D-GRE sequence with a relatively long repetition time (TR) and short echo time (TE).^9^ For COSMOS QSM reconstruction, a 3D-GRE sequence is also utilized to acquire data at three or more head orientations with a relatively long TE for improved phase contrast.^23^ Herein, we exploit the unused TR time in the MTV acquisition to collect additional late echoes for QSM processing. Additionally, we collect a different FA at each head orientation, allowing for simultaneous MTV and COSMOS reconstruction without scan time penalty.

To further accelerate the acquisition, we employ JVC-GRAPPA which jointly reconstructs multi-echo images by treating data from other echoes as extra coils, improving parallel imaging capability.^17^ The VC concept is applied to double the number of channels. The k-space signal received from a coil can be represented by the spatial spin-density distribution weighted by the spatial complex coil s’ensitivity.^21^ The symmetric complex-conjugate signal can be interpreted as the signal received by a virtual coil. While the magnitude sensitivity from the virtual coil is the same as the actual coil, the phase is different, providing additional encoding power. As each coil receives contributions from all coils and all echoes, for N_c_ coils and N_e_ echoes, we would train (2 x N_c_ x N_e_)^2^ kernels.

To address the problem that arises from the large number of kernels that need to be estimated (proportional to square of the number of channels), an iterative procedure is performed with an initial joint GRAPPA reconstruction. The entire k-space data estimated using the initial joint GRAPPA are then utilized in training successive JVC-GRAPPA kernels^19^ Additionally, k-space sampling patterns of the individual echoes are shifted with respect to each other, providing complementary frequency information.

### 2.1. Data Acquisition

Three healthy adult volunteers were scanned on a 3T scanner (Siemens Skyra) with a 64-channel head coil in accordance with IRB-approved protocol.^24^

#### Subject #1

3D multi-echo GRE sequence was acquired at three head orientations (0°, 7°, and 13° with respect to B_0_; the same approximate orientations were also acquired for Subjects #2-4) with the following parameters: 3 echoes, TE_1_=5.73 ms, echo spacing=9.14 ms, TR=31 ms, FOV=192×256×160 mm^3^, and the vendor’s R=2×2 GRAPPA. A different FA (4°, 10°, 20°) was applied at each head orientation. Rapid B_1_+ mapping calibration was acquired at each orientation using a 3D-FLASH sequence (acquisition time = 31 seconds at each orientation).^25^ The total acquisition time was approximately 15 min and 17 min with B1+ mapping. Additional *single orientation* data was separately acquired.

#### Subject #2

Fully sampled data were acquired at three head orientations (acquisition time = 60 min; 62 min with B1+), from which undersampling was *retrospectively* performed.

#### Subject #3

*Prospective* undersampling was performed including R=3×2 GRAPPA (acquisition time = 9 min; 11 min with B1+) and R=3×3 JVC-GRAPPA reconstruction (acquisition time = 6 min; 8 min with B1+). The latter was enabled by introducing gradient blips to the sequence to produce k-space shifting at later echoes.

#### Subject #4

To evaluate the reproducibility of multi-orientation MTV, scan-rescan was performed using the imaging same parameters as Subject #1. This included two sets of multi-orientation and single orientation 3D multi-echo GRE data.

### 2.2. Data Processing

Subject #1’s data were reconstructed with the vendor’s online GRAPPA. From subject #2’s fully-sampled data, undersampling was performed retrospectively along two phase-encoding axes by acceleration factors of R=3×2 for GRAPPA and R=3×3 for JVC-GRAPPA. An ACS region of size 24×24 and a kernel size of 3×3 were used for both methods. During reconstruction, Tikhonov-regularization parameters were used for kernel calibration of R=3×2 GRAPPA (λ=10^-8^) and R=3×3 JVC-GRAPPA (λ=10^-7^) to optimize for root mean squared error (RMSE) relative to the fully-sampled ground truth. For subject #3, in order to provide complementary k-space information, the sampling pattern was shifted by (1,1) in the second echo, (2,2) in the third echo, and (3,3) in the fourth echo relative to the first echo.

For *susceptibility*, the raw phase data were processed using Laplacian-based phase unwrapping.^26^ Background phase removal was performed using V-SHARP in STI-Suite with a maximum kernel size of 25 voxels. The tissue phase images were then registered to the neutral orientation using FSL-FLIRT (https://fsl.fmrib.ox.ac.uk). COSMOS reconstruction was performed using an iterative least-squares formulation.^14^

For *MTV*, the B_1_+ map at each head orientation was co-registered to the corresponding GRE magnitude image using FSL-FLIRT. Calculation of the effective FA for B1+ bias-correction at each orientation was performed using the double-flip angle method.^12,27^ The bias-corrected magnitude images were then skull-stripped and registered to the neutral orientation. T_1_ and M_0_ (product of the coil reception profile and proton density [PD]) maps were calculated using weighted-least-squares fitting.^9^ The reception profile was estimated using a method based the linear relationship between PD and T1 as previously described.^28,29^ PD maps were subsequently normalized to CSF-only voxels (WF), from which MTV was calculated.

R_2_*-mapping was performed using a least-squares mono-exponential fit of the multi-echo magnitude data. The lsqnonneg function in Matlab (v.R2015b) was used to apply a nonnegative constraint to generate realistic values.

### 2.3. Data Analysis

To compare multi-orientation to single-orientation MTV in subject #1, we coregistered the multi-orientation GRE acquisitions to the neutral frame using *niftyreg* (http://cmictig.cs.ucl.ac.uk/wiki/index.php/NiftyReg) after correcting for B_1_+ inhomogeneity. The co-registered images were compared to their single-orientation counterparts using RMSE. This was also performed for the MTV maps (multi-versus single-orientation). In addition, regions of interests (ROIs) were manually placed by a board-certified neuroradiologist (F.Y., 8 years of experience) within selected brain regions using 3D Slicer (v.4.10.2, https://www.slicer.org/).^23,30^ Pearson correlation was used to measure the correlation between the single-versus multi-orientation MTV ROIs.

For subject #2’s *retrospectively* undersampled data, RMSE was calculated for R=3×2 GRAPPA and R=3×3 JVC-GRAPPA relative to fully sampled data. To assess similarity of *prospectively* undersampled JVC-GRAPPA to GRAPPA in subject #3, RMSE was computed between the two sets of reconstructed images.

### 2.4. Contribution of myelin and iron to susceptibility

A potential application of the multiparametric maps is in separating the contributions of iron and myelin to susceptibility. We computed iron-induced and myelin-induced susceptibility maps based on the SEMI-TWInS method.^31^ SEMI-TWInS employs a three-compartment tissue model composed of iron (Fe), myelin (My), and extracellular matrix (M). Two key assumptions are made about the relationship between R_2_*, bulk susceptibility *χ*, iron concentration *C_Fe_*, and myelin concentration *C_My_*. The first states that R_2_* and *χ* can be derived using linear combinations of *C_Fe_* and *C_My_*, wherein

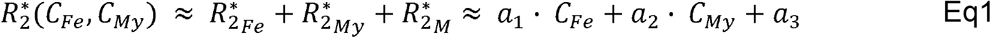

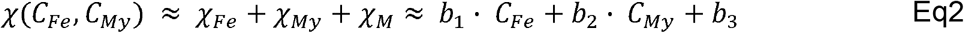

The second assumption states that each coefficient (*a*_1_, *a*_2_, *a*_3_, *b*_1_, *b*_2_, *b*_3_) maintains a constant value regardless of brain tissue type, and is unique to each individual. Of note, *C_Fe_* and *C_My_* represent spatially varying quantitative maps while *a*_1_, *a*_2_, *a*_3_, *b*_1_, *b*_2_, *b*_3_ are constants.

The six coefficients were computed by first determining R_2_*, *χ*, *C_Fe_*, and *C_My_* and subsequently solving Equations 1 and 2 using the NumPy linear least-squares solver. ROIs were manually drawn in ten brain regions. Within the ROIs, R_2_*, *χ*, and *C_My_* (MTV) were measured on a voxel-wise basis while *C_Fe_* was taken from literature values.^32^ The coefficients were computed by consolidating R_2_*, *χ*, *C_Fe_*, and *C_My_*, and then least-squares solving Eqs. (1) and (2).

was then computed in the brain by solving Equations 1 and 2 with respect to R_2_*,, and the estimated coefficient values (). Iron-induced susceptibility and myelin-induced susceptibility maps were computed as and, respectively.

## 3. Results

### 3.1. Single versus multi-orientation

Previous studies used a single head orientation to estimate MTV, whereas we implemented a multiple head orientations approach.^9,16^ The multi-orientation B_1_+ corrected magnitude data had RMSEs of 4.8-5.8% relative to single orientation data (the same neutral orientation FA=20° data was used for multi- and single-orientation). Multi-orientation MTV demonstrated an RMSE of 4.5% compared to single-orientation MTV (Figure 1), with the two maps being strongly correlated (Supporting Information Figures S1 and S2). Additional parametric maps (T_1_, PD, R_2_*) were generated without added scan time (Figure 2).

**Figure 1:**
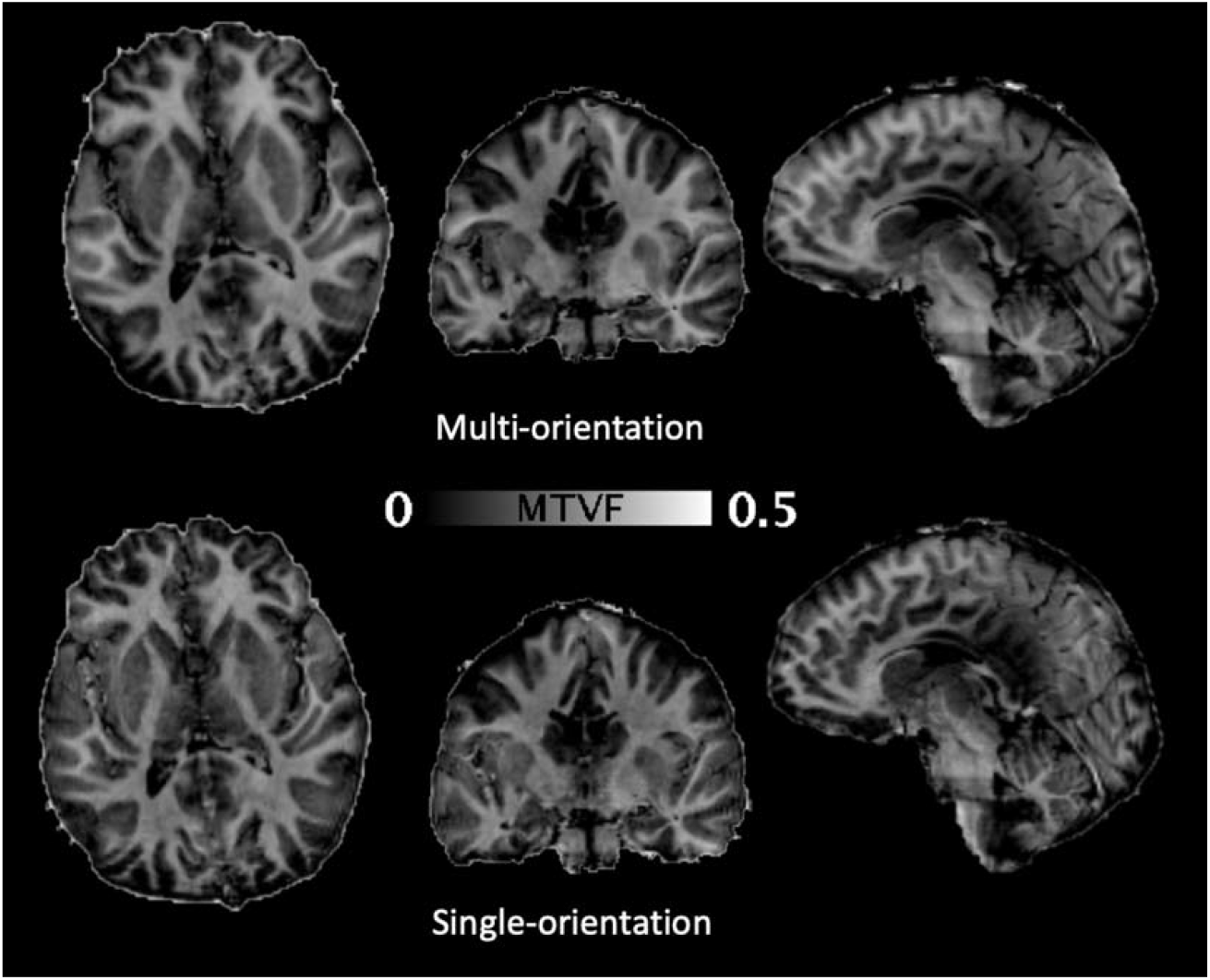
MTV maps acquired using single-(bottom row) and multiple-head orientations (bottom row) for Subject #1 presented in axial, coronal, and sagittal projections, which demonstrate similar results.

**Figure 2:**
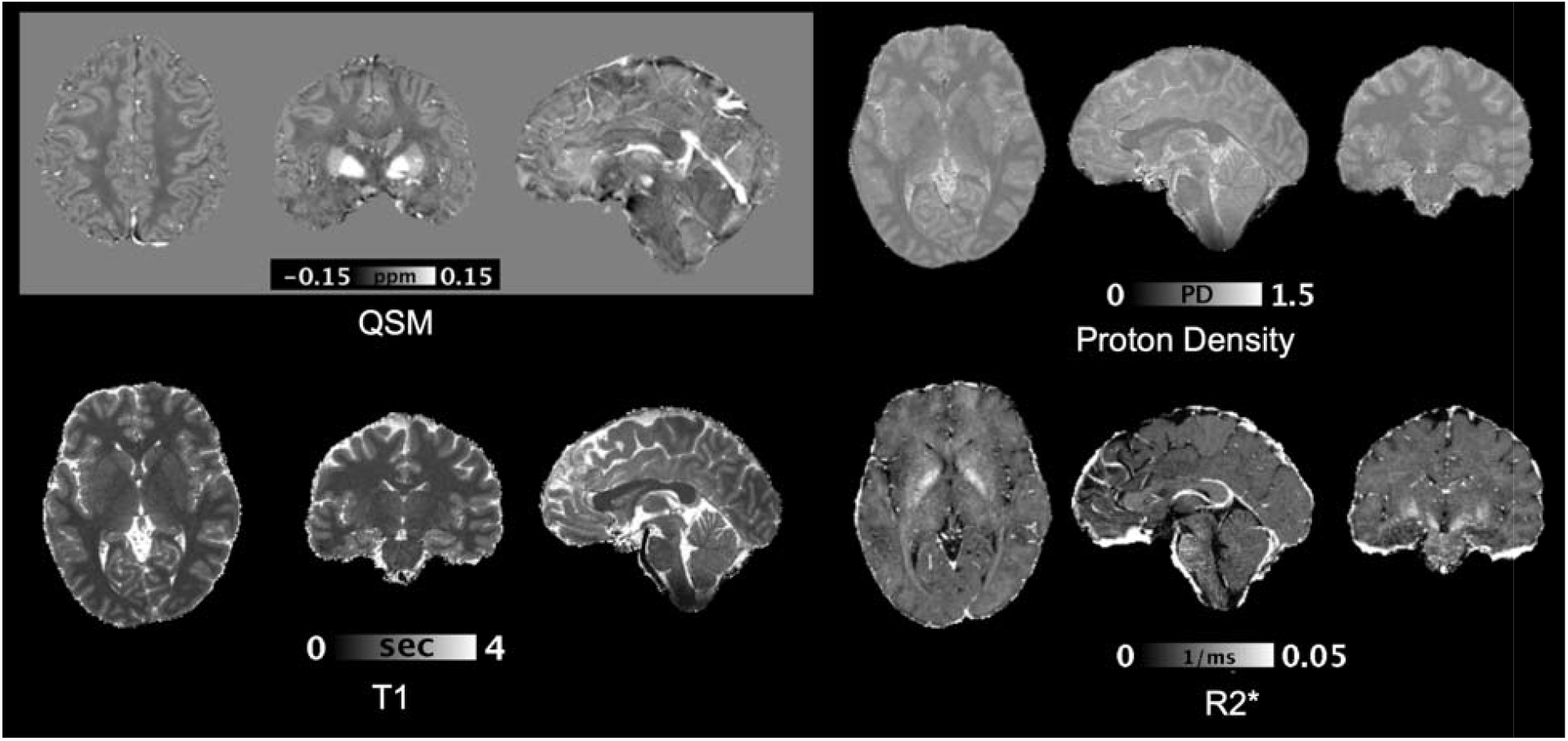
Representative multi-parametric maps (QSM, R2*, T1, and PD) generated from multi-orientation data for Subject #2.

Scan-rescan data for Subject #4 demonstrated an RMSE of 3% between the initial and the repeat multi-orientation MTV maps, as compared to an RMSE of 3.2% between initial set of multi-orientation and single orientation MTV maps (Supporting Information Figure S3). The single orientation data demonstrated an RMSE of 1.8% between the initial and repeat single-orientation MTV data.

### 3.2. Fully sampled versus retrospectively undersampled data

For FA=4°, R=3×2 GRAPPA and R=3×3 JVC-GRAPPA yielded RMSEs of 3.1% and 3.5% with respect to R=1 data. For FA=10°, GRAPPA and JVC-GRAPPA yielded RMSEs of 4.2% and 4.7%, respectively. RMSEs for FA=20° were 5.3% and 5.7% for GRAPPA and JVC-GRAPPA, respectively. Comparing retrospectively undersampled MTV maps to R=1, GRAPPA and JVC-GRAPPA demonstrated RMSEs of 7.5% and 7.7% (Supporting Information Figures S4 and S5).

### 3.3. Prospectively acquired undersampled data

Comparing JVC-GRAPPA magnitude images to GRAPPA, RMSEs for FA=4°, 10°, and 20° were 3.1%, 5.1%, and 6.5%, respectively. Moreover, JVC-GRAPPA retained higher SNR, particularly for later echoes (Figure 3).

**Figure 3:**
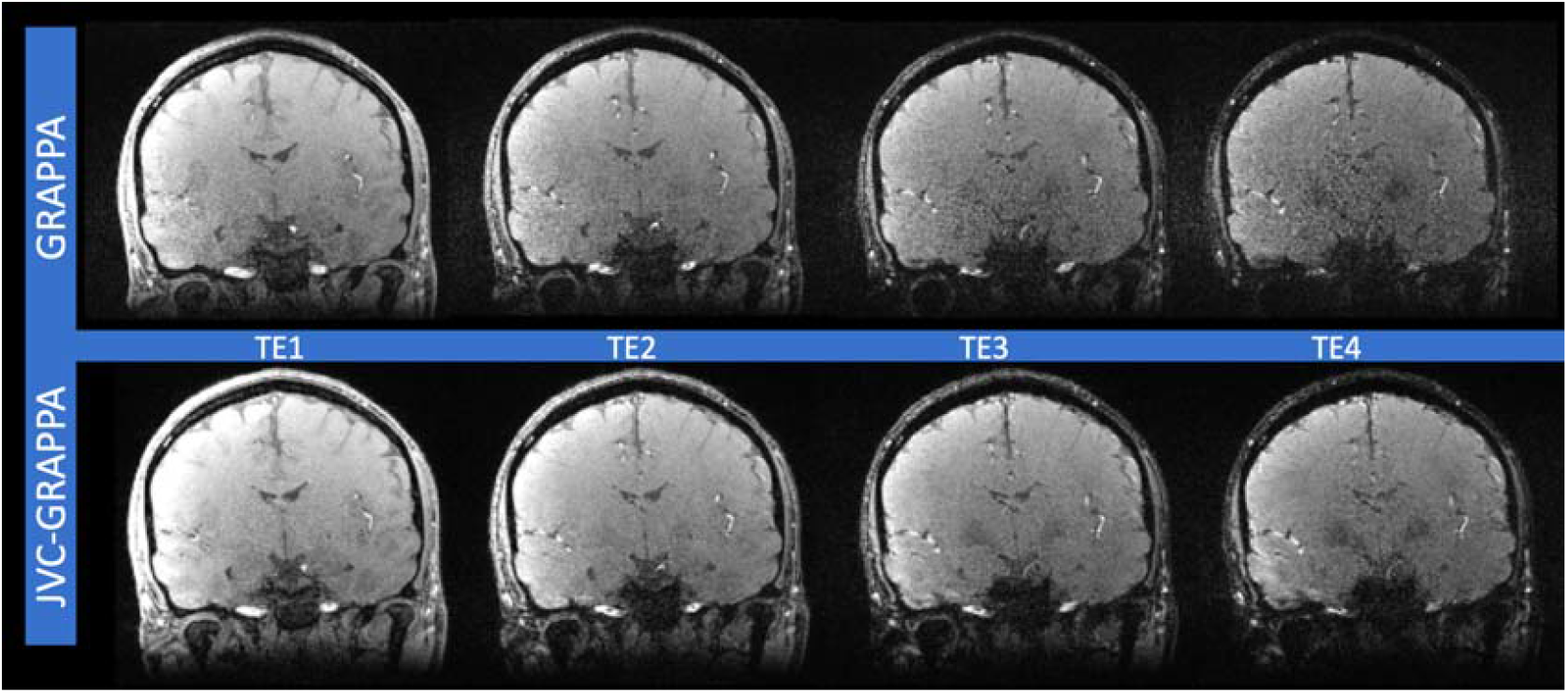
Prospectively under-sampled multi-echo GRE magnitude images obtained using R=6 GRAPPA (top row) and R=9 JVC GRAPPA (bottom row) from Subject #3. JVC-GRAPPA reconstruction demonstrates higher SNR particularly for late echoes.

QSM demonstrated an RMSE of 1.4% between JVC-GRAPPA and GRAPPA, with no significant differences in susceptibility values (p>0.1; Supporting Information Figures S6 and S7). Comparing JVC-GRAPPA and GRAPPA reconstruction-based MTV yielded an RMSE of 5.9% (Figure 4; Supporting Information Figures S8 and S9), with larger standard deviations observed with GRAPPA (p=0.002). We note that in the absence of ground truth, RMSEs computed between JVC-GRAPPA and GRAPPA aim to measure their similarity.

**Figure 4:**
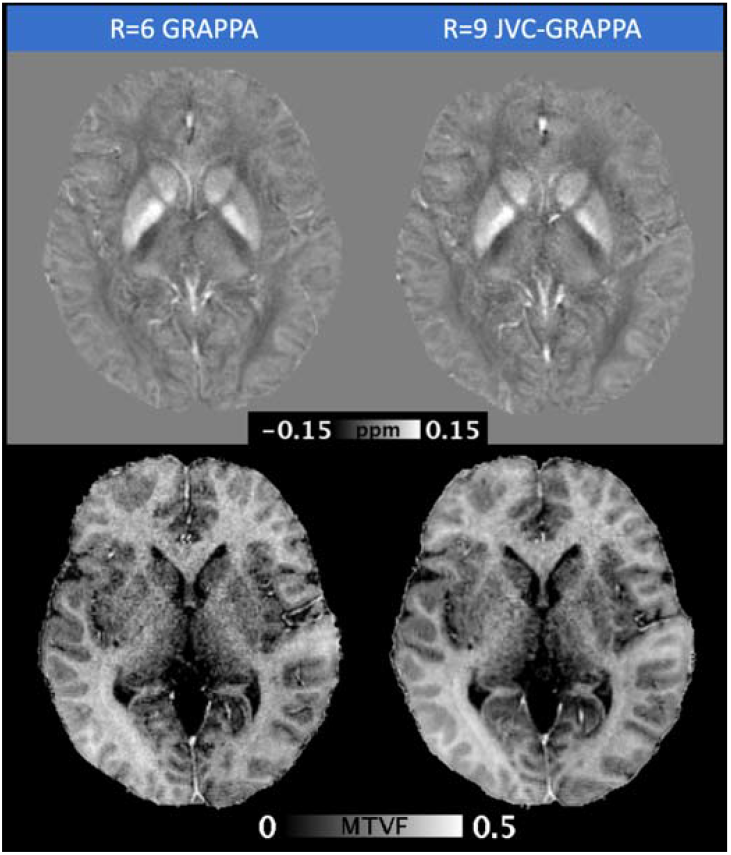
QSM (top) and MTV (bottom) maps generated from prospectively undersampled R=6 GRAPPA (left) and R=9 JVC-GRAPPA (right) reconstructions from Subject #3. No significant differences can be observed with visual inspection.

### 3.4. Myelin and iron contributions to susceptibility

Linear least-squares fitting yielded the following coefficients: = 0.001 Hz*100mg/mg; = 0.013 Hz/MTV; = 0.012 Hz; = 0.009 ppm*g/mg; = −0.040 ppm/MTV; = −0.022 ppm. The map demonstrated significant differences from (Figure 5). The hypointense signal within the map corresponds to negative susceptibility values, reflecting diamagnetic myelin. Image contrast was particularly striking between grey and white matter. Within deep gray nuclei, myelin-induced susceptibility contributions could be observed, although the iron-related contribution was predominant.

**Figure 5:**
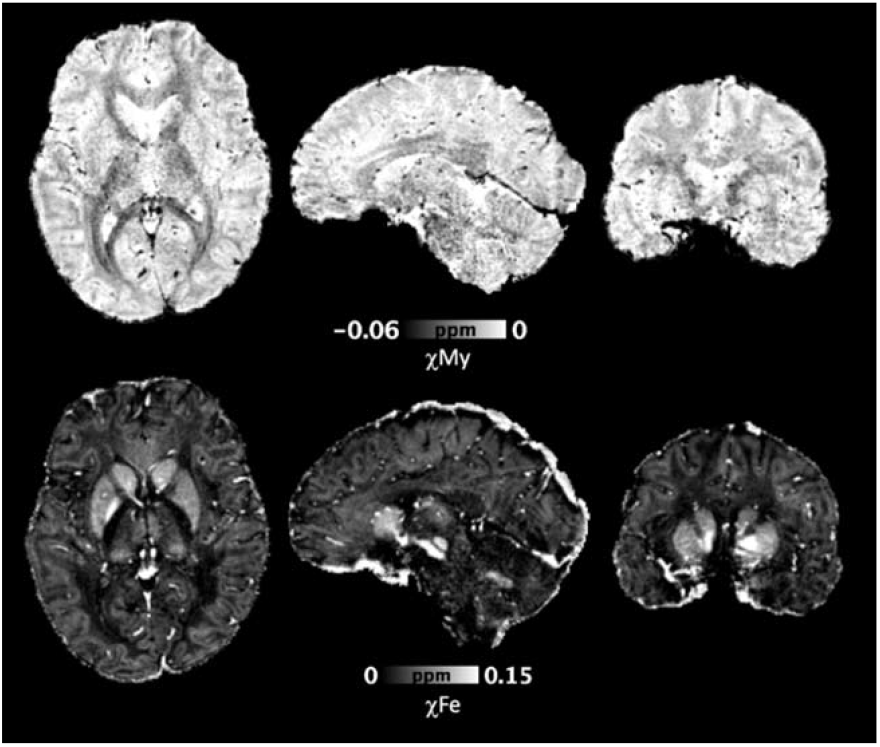
Representative myelin (χMy) and iron (χFe)-induced susceptibility maps presented in axial, sagittal, and coronal projections. Diamagnetic signal is present within the white matter, but also the subcortical grey matter, consistent with the known distribution of myelin within the brain. Paramagnetic signal is most pronounced within the subcortical grey matter, but also seen within the cerebral cortex and to a lesser degree within the white matter.

## 4. Discussion

We propose an acquisition strategy that leverages inherent redundancies to obtain MTV and QSM within the same scan time as one parametric map, permitting concurrent interrogation of myelin content and magnetic susceptibility. We were also able to obtain three additional parametric maps (PD, T_1_, R_2_*) without scan time penalty. By applying JVC-GRAPPA reconstruction, we reduced the total acquisition time to 6 minutes.

### 4.1. JVC-GRAPPA

Our multi-echo acquisition supports the use of joint reconstruction, which was leveraged to decrease scan time, making the proposed method more suitable for clinical use. The shifted undersampling patterns of k-space across later echoes allowed for increased frequency coverage. In addition, VC concept was utilized to double the number of available channels to enhance spatial encoding. While there was noise amplification resulting from the intrinsic SNR penalty associated with parallel imaging, the resulting images had improved SNR compared to the slower GRAPPA.

One potential limitation is that because the method stacks data from all image contrasts along the coil axis, larger amounts of calibration data are required. The shifted k-space sampling further contributes to this as the staggered JVC kernels span a larger k-space extent, expanding the ACS region.^17^ To alleviate this, we used an initial joint GRAPPA reconstruction to provide an estimate of the entire k-space matrix, which was used for calibrating the following JVC steps. Compared to other highly-accelerated methods, advantages of JVC-GRAPPA include its relative ease in processing and lower computational demands, which facilitate potential online reconstruction.^17^

### 4.2. MTV

Previous studies have acquired multi-FA GRE images in a single head orientation.^9^ In this study, after performing B_1_+inhomogeneity correction for each orientation, the RMSE between the magnitude images was minimal.^33^ Although GRE magnitude images are influenced by orientation of tissues relative to B_0_, we found that the resultant multi- and single-orientation MTV maps yielded similar results.^30^

The GRE images demonstrated minimal differences between R=9 JVC-GRAPPA and R=6 GRAPPA reconstructions. MTV demonstrated a slightly greater difference between acceleration methods, which likely reflects additional noise amplification as a result of the intervening processing steps.

### 4.3. QSM

COSMOS reconstruction has been shown to provide higher quality susceptibility estimates than regularized single orientation methods.^6,14^ Generally, single orientation reconstructions can suffer from streaking artifacts as well as over-smoothing. However, in order to achieve optimal imaging using COSMOS, sampling at a minimum of three head orientations is required.^23^ Significant deviations, either through fewer than three acquisitions or reducing the angle between sampling orientations, can result in streaking artifacts.

We acknowledge that even with reduced scan time, there are patients for whom adequately rotating the head may prove challenging. In these circumstances, the proposed method can be acquired using a single head orientation, with subsequent application of a QSM regularization method.^6,34^ Although the estimation within anisotropic tissues could vary depending on head orientation due to susceptibility anisotropy^35^, isotropic structures or those with greater intrinsic susceptibility (deep grey nuclei) should appear similar to COSMOS.^30,34^

### 4.4. Additional contrasts

R_2_*, T_1_, and PD maps were also generated from the acquired data without additional scan time cost. R_2_* is influenced by myelin and iron content.^36^ While R_2_* can be calculated from a single orientation, combining all three yielded higher SNR. T_1_-mapping has potential applications in the evaluation of hepatic encephalopathy^37^ and multi-system atrophy^38^. Recently, Filo et al. used a combination of parametric measures, including T1 (R1) and MTV, to reveal region-specific molecular changes associated with aging.^39^ A similar strategy could be applied to detect pathologic deviations from normal aging that occur in neurologic diseases.

We showed that by leveraging the intrinsically co-registered multi-parametric maps (R_2_*, susceptibility, MTV), high-spatial-resolution parametric maps of iron and myelin-induced susceptibility can be generated within the same anatomic space. These maps are relevant because although studies have supported using QSM as a brain iron marker, myelin also affects susceptibility particularly within white matter^40^. It is important to acknowledge that MTV is not entirely specific for myelin and is affected by other diamagnetic macromolecules^9^. Likewise, in certain brain regions, other non-iron paramagnetic substrates could influence susceptibility. Nonetheless, the ability to approximate the voxel-wise contributions of myelin and iron to susceptibility enhances the clinical utility of magnetic susceptibility imaging.^41^ Potential applications include demyelinating diseases, traumatic brain injury, and neurodegenerative diseases.

### 4.5. Limitations

Joint parallel imaging is associated with increased reconstruction time compared to GRAPPA.^17^ Applying the JVC kernels using a less demanding element-wise multiplication in image space (rather than convolution in k-space) could help address this. To delineate susceptibility contributions from iron, we utilized literature reference values for iron concentration for calibration. We also note that paramagnetic and diamagnetic substrates other than myelin and iron may contribute to the observed susceptibility. Future plans include exploring alternative calibration schemes that may allow for more robust estimations in neuropathology. A current limitation is our small study size, restricting generalizability of the results. To realize the potential of our quantitative imaging approach, it will be necessary to image additional subjects and across different sites.^42^

## 5. Conclusion

In this study, we proposed a strategy for simultaneously acquiring QSM and MTV by exploiting their inherent redundancies to reduce scan time. We also generated multiple quantitative parametric maps, including T_1_, R_2_*, *χ_My_*, *χ_Fe_*, and PD without scan time penalty. Clinical feasibility was enhanced by further accelerating the acquisition by 9-fold using JVC-GRAPPA while preserving image quality. This protocol has potential applications in the quantitative evaluation of neurological diseases.

## Figure Captions

Supporting Information Figure S1: Selected grey and white matter regions in the brain (Subject #1) with their corresponding values on single and multi-orientation MTV (□=0.965, p<0.0001).

Supporting Information Figure S2: 2D histogram of voxel-wise correlation over the whole brain between single and multi-orientation MTV for Subject #1.

Supporting Information Figure S3: Scan-Rescan axial MTV images from Subject #4 (from left to right: initial multi-orientation MTV, repeat multi-orientation MTV, and initial single-orientation MTV).

Supporting Information Figure S4: 2D histogram of voxel-wise correlation over the whole brain between fully sampled R=1 and retrospectively undersampled R=6 GRAPPA MTV for Subject #2.

Supporting Information Figure S5: 2D histogram of voxel-wise correlation over the whole brain between fully sampled R=1 and retrospectively undersampled R=9 JVC-GRAPPA MTV for Subject #2.

Supporting Information Figure S6: Selected grey and white matter regions with their corresponding values on R=6 GRAPPA and R=9 JVC-GRAPPA for QSM, respectively.

Supporting Information Figure S7: 2D histogram of voxel-wise correlation over the whole brain between prospectively undersampled R=6 GRAPPA and R=9 JVC-GRAPPA QSM for Subject #3.

Supporting Information Figure S8: Selected grey and white matter regions with their corresponding values on R=6 GRAPPA and R=9 JVC-GRAPPA for MTV, respectively.

Supporting Information Figure S9: 2D histogram of voxel-wise correlation over the whole brain between prospectively undersampled R=6 GRAPPA and R=9 JVC-GRAPPA MTV for Subject #3.

